# *In vivo* CRISPR screens reveal SCAF1 and USP15 as novel drivers of pancreatic cancer

**DOI:** 10.1101/2022.03.13.484174

**Authors:** Sebastien Martinez, Ramona Weber, Tristan Woo, Ahmad Malik, Michael Geuenich, Gun Ho Jang, Dzana Dervovic, Khalid N. Al-Zahrani, Ricky Tsai, Nassima Fodil, Philippe Gros, Sachdev S Sidhu, Stephen Gallinger, G. Gregory Neely, Kieran Campbell, Faiyaz Notta, Ataman Sendoel, Daniel Schramek

## Abstract

Functionally characterizing the genetic alterations that drive pancreatic cancer progression is a prerequisite for Precision Medicine. Here, we developed a somatic CRISPR/Cas9 mutagenesis screen to assess the transforming potential of 125 recurrently mutated ‘long-tail’ pancreatic cancer genes, which revealed USP15 and SCAF1 as novel and potent Pancreatic ductal adenocarcinoma PDAC tumor suppressors, with USP15 functioning in a haplo-insufficient manner. Mechanistically, we found that loss of USP15 leads to reduced inflammatory responses associated with TNFα, TGF-β and IL6 signaling and sensitizes pancreatic cancer cells to PARP inhibition and gemcitabine. Similarly, genetic ablation of SCAF1 reduced inflammatory responses linked to TNFα, TGF-β and mTOR signaling and increased sensitivity to PARP inhibition. Furthermore, we identified that loss of SCAF1 resulted in the formation of a truncated inactive USP15 isoform at the expense of full length USP15, functionally coupling SACF1 and USP15. Notably, USP15 and SCAF1 mutations or copy number losses are observed in 31% of PDAC patients. Together, our results demonstrate the utility of *in vivo* CRISPR to integrate human cancer genomics with mouse modeling to delineate novel cancer driver genes *USP15* and *SCAF1* such as with potential prognostic and therapeutic implications.

## Introduction

Pancreatic ductal adenocarcinoma (PDAC) is the fourth-leading cause of cancer-related death in industrialized countries and is predicted to be the second-leading cause of cancer death in the United States by 2040^1,2^. Despite recent progress in our understanding of the molecular and genetic basis of this pathology, 5-year survival rates remain low and do not exceed 10%. PDAC is an epithelial tumor that arises from the cells of the pancreatic duct and represents the vast majority of pancreatic neoplasms. PDAC develops due to the acquisition of cooperating alterations in tumor suppressor and oncogenes as well as chromosomal aberrations, which are thought to either occur gradually by a multi-step process or simultaneously in a single catastrophic event^3,4^. Through these mutational processes tumors also accumulate hundreds of random bystander mutations, which make it exceedingly hard to interpret genomic data and identify the few real driver mutations that trigger tumor initiation, progression, metastasis and therapy resistance. Whole exome sequencing studies identified a number of frequent mutations altering the function of key oncogenes and tumor suppressor such as *KRAS* (93%), *TP53* (72%), *CDKN2A* (44%), *SMAD4* (40%), *RNF43* (8%), *FBXW7* (5%)^5–7^.

In the clinic, genomic technologies are reaching the point of detecting genetic variations at high accuracy in patients. This holds the promise of fundamentally alter clinical practice by personalizing treatment decisions based on the genetic make-up of an individual tumor, commonly referred to as Precision Medicine. These genomic advances have validated previous findings regarding the most commonly mutated PDAC genes, but also led to the identification of a long-tail of recurrent but less frequent alterations in hundreds of genes^8,9^. Multiple hypotheses have been proposed to explain the low frequency and high diversity of those infrequently mutated genes. Some of those mutations might provide a functional alteration similar to that of a major driver. Some long tail mutations may as well be highly penetrant but simply affect genes that are rarely mutated^10^. Alternatively, some might affect the same pathway or molecular mechanism and cooperate to promote tumor progression as recently showed by our study of rarely mutated long-tail genes in head and neck cancer converging on inactivation of the NOTCH signaling pathway^11^. These long-tail genes often lack biological or clinical validation and their contribution to PDAC development remains unknown. As such, establishing, reliable and genetically traceable *in vivo* screening platforms to systematically identify putative PDAC driver genes, is a prerequisite to fulfill the promise of Precision Medicine.

Genetically engineered mouse models (GEMM) constitute the gold standard for genetic perturbation studies. Mouse models of human cancers have provided invaluable insights into the genes and molecular mechanisms that drive cancer development^12,13^ and proven essential as preclinical model in the development of novel therapeutic agents^14^. However, conventional GEMMs are extremely time and resource intensive rendering them inapt to sift through the scores of genetic alterations emerging from large-scale genomics projects^15^. Here, we report an *in vivo* CRISPR/Cas9 screening strategy to identify which long-tail PDAC genes and associated pathways cooperate with the oncogenic *Kras^G12D^* to accelerate pancreatic cancer progression.

## RESULTS

### Direct *in vivo* CRISPR gene editing in the mouse pancreas

To functionally test putative PDAC cancer genes *in vivo*, we employed a multiplexed CRISPR/Cas9 genome editing approach to generate knock-out clones directly in pancreatic epithelium of tumor-prone mice. We used conditional Lox-Stop-Lox-(LSL)-*Kras*^G12D^ and a LSL-*Cas9-GFP* mice crossed to the pancreas-specific PDX1-Cre driver line (termed KC mice) and injected an adeno-associated virus that expresses an sgRNA and the H2B-RFP fluorescent marker (AAV-sgRNA-RFP) **(Fig. 1a)**. Cre-mediated excision of Lox-Stop-Lox cassettes resulted in expression of oncogenic *Kras^G12D^, Cas9* and *GFP* and formation of hundreds of cytokeratin19 positive (CK19) pancreatic intraepithelial neoplasia (PanIN) precursor lesions, which can be lineage-traced by virtue of red fluorescence (**Supplementary Fig. 1a, b**). To validate the efficiency of CRSIPR/Cas9-mediated mutagenesis, we injected sgRNAs targeting *GFP*, which revealed a knock-out efficacy of 78±6% (**Supplementary Data Fig. 1c**).

**Fig. 1.**
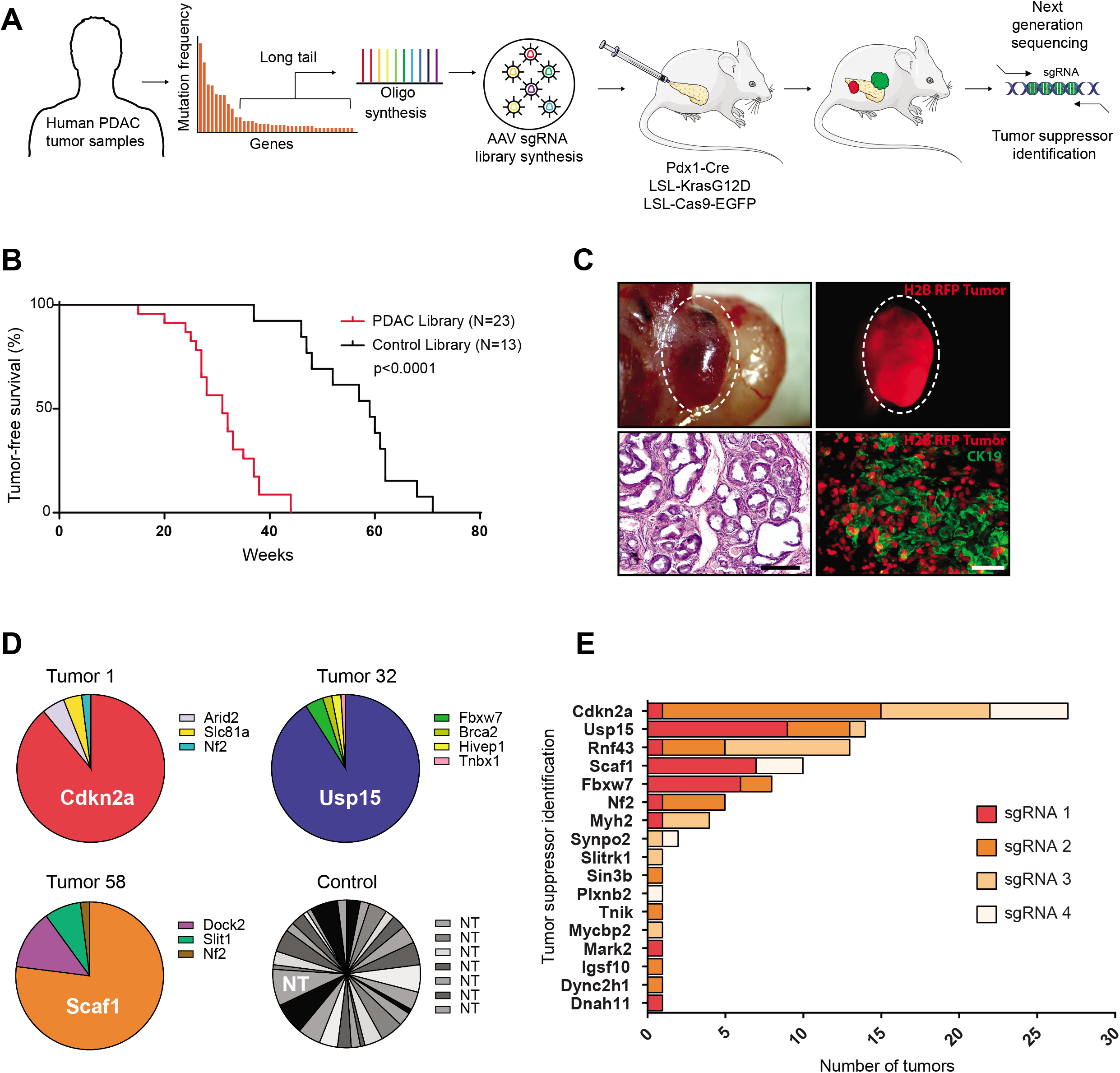
*In vivo* CRISPR screen reveals novel pancreatic cancer tumors suppressors. **A**, Experimental design of the *in vivo* PDAC CRISPR screen, showing gene selection from long-tail mutations, pancreatic injection of AAV libraries and tumor sequencing. **B**, Tumor-free survival of Pdx1-Cre;LSL-*Kras*^G12D^;LSL– *Cas9-GFP* mice transduced with a sgRNA library targeting putative pancreatic cancer genes or a control sgRNA library. **C**, Representative images of an H2B-RFP+ pancreatic PDAC-library tumor. Representative H&E images showing a PDAC-library pancreatic tumor. Scale bar 250μm. Representative immunofluorescence of PDAC-library tumor showing H2B-RFP and CK19 expression. Scale bar 50 μm. **D**, Representative pie charts showing tumor suppressor genes with enriched sgRNAs in tumor DNA obtained from three different pancreatic tumors and a control-transduced pancreas with multifocal PanINs. **E**, Bar graph showing putative tumor suppressor genes with enriched sgRNAs in tumor DNA obtained from the PDAC mouse model (sgRNA enriched per tumors are indicated by color).

KC mice exhibited rapid growth of pre-invasive PanINs precursor lesions but they show a very slow progression to invasive PDAC with a median latency of 14 month (**Fig. 1b**). Additional genetic alterations such as loss of transformation related protein 53 *(Trp53), p16Ink4a, Lkb1* or inactivation of TGF-β signaling was previously shown to cooperate with *Kras*^G12D^ and induces rapid PDAC development within 3-5 month^16–21^. To test whether our direct *in vivo* CRISPR approach can reveal genetic interactions, we recapitulated cooperation between oncogenic *Kras*^G12D^ and loss of p53 (*Trp53*). Indeed, Cas9-mediated ablation of *Trp53* in KC mice triggered rapid PDAC formation with a median latency of 14 weeks, while littermates transduced with scrambled control sgRNAs remained cancer-free for over 1-year (**Supplementary Data Fig. 1d**). This is in line with previous efforts using CRISPR/Cas9 gene editing in *KRas*^G12D^ mice ^22,23^ and demonstrates that this approach can be used to test for genetic cooperation between PDAC genes.

### CRISPR Screen identifies novel PDAC tumor supressors

In pancreatic cancer, 125 genes show recurrent somatic mutations^6,7^. To assess these genes *in vivo*, we established a sgRNA library targeting the corresponding mouse orthologs (4 sgRNAs/gene; 500 sgRNAs) as well as a library of 420 non-targeting control sgRNAs (**Supplementary Table 1**). Of note, we did not include sgRNAs targeting well-established PDAC driver genes such as *Trp53* or *Smad4*^16–21^.

Next, we optimized the parameters for an *in vivo* CRISPR screen. Using a mixture of AAV-GFP and AAV-RFP we determined the viral titer that transduces the pancreatic epithelium at clonal density (MOI<1). Higher viral titers were associated with double infections, whereas a 15% overall transduction level minimized double infections while generating necessary clones to screen (**Supplementary Data Fig. 1e**). Using multicolor Rosa26-Lox-Stop-Lox(R26-LSL)-Confetti Cre-reporter mice, we next determined the viral titer required to generate thousands of discrete clones within the pancreatic epithelium (**Supplementary Data Fig. 1f**). Thus, at a transduction level of 15% and a pool of 500 sgRNAs, each sgRNA would be introduced into at least 50 CK19+ epithelial cells within a single pancreas.

To uncover long-tail genes that cooperate with oncogenic *KRas^G12D^* and accelerate PDAC development, we injected the experimental and the control AAV-sgRNA libraries into the pancreas of 23 and 13 KC mice, respectively. Next generation sequencing confirmed efficient AAV transduction of all sgRNAs (**Supplementary Data Fig. 2a**). Importantly, KC mice transduced with the long-tail PDAC sgRNA library developed pancreatic cancer significantly faster than littermates transduced with the control sgRNA library (31 versus 59 weeks; p<0.0001) (**Fig. 1b and c**). In addition, 13/23 (56%) KC mice transduced with the long-tail PDAC sgRNA library developed liver and/or lung metastasis, while only 1/13 (~8%) littermate mice transduced with the control sgRNA library developed metastasis (**Supplementary Data Fig. 2b, c**), indicating the existence of strong tumor suppressors within the long-tail of PDAC associated genes.

To identify these PDAC driver genes, we examined the sgRNA representation in 151 tumors. 78% of tumors showed strong enrichment for a single sgRNAs, indicating a clonal origin. In contrast, the pancreas of control-transduced mice with multifocal PanINs showed enrichment of several non-template control sgRNAs (**Fig. 2d**). We prioritized genes that were targeted by ≥2 sgRNAs and knocked-out in multiple tumors, resulting in 8 candidate tumor suppressor genes (**Fig. 1e, Supplementary Table 2**). These candidates included well-known PDAC tumor suppressor genes, such as *Cdkn2a^24^, Rnf43^25^, Fbxw7^26^* or *NF2^27^*, as well as genes with poorly understood function, such as *Usp15* and *Scaf1*.

**Fig. 2.**
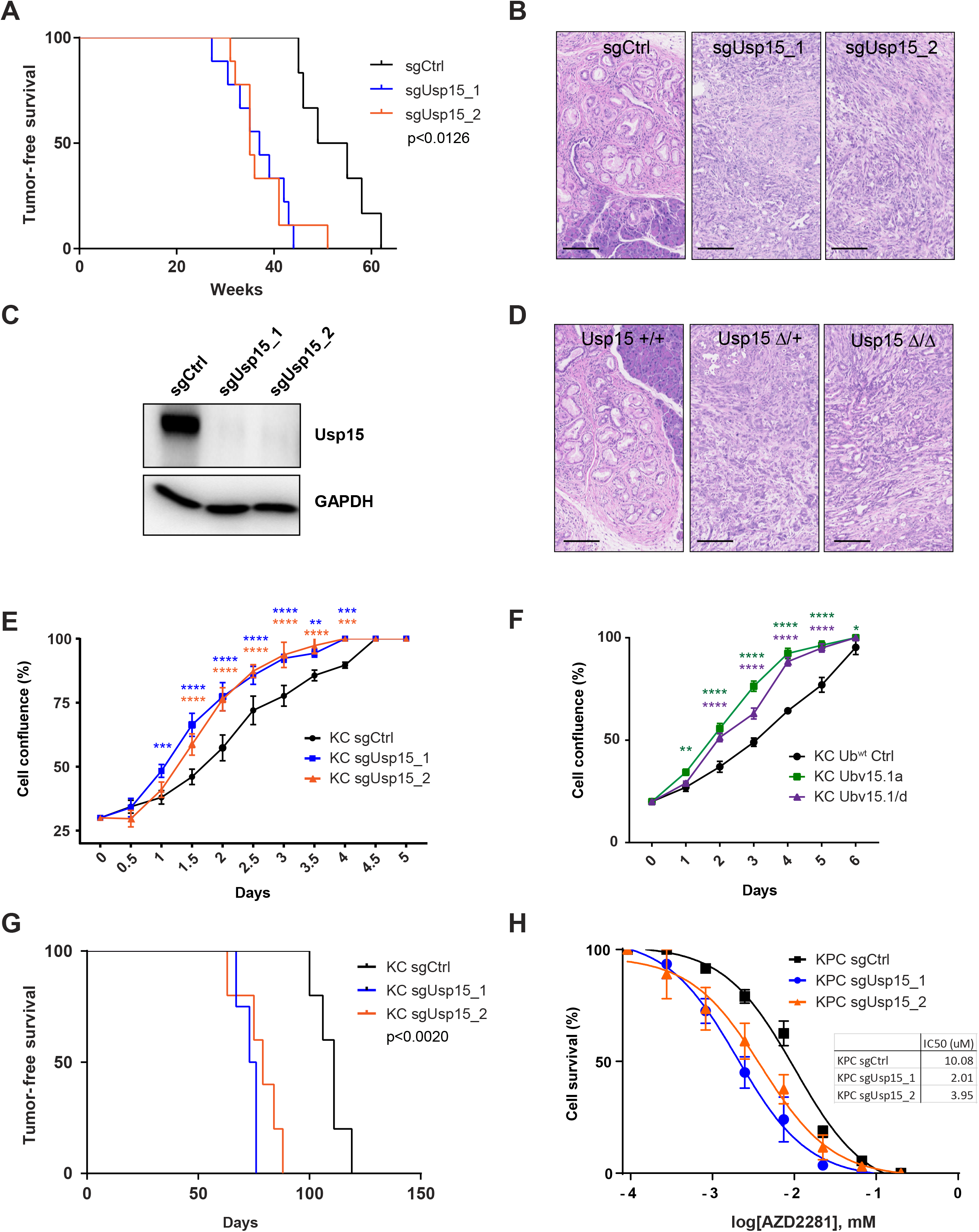
Usp15 functions as PDAC tumor suppressor. **A,** Tumor-free survival of Pdx1-Cre;LSL-*Kras*^G12D^;LSL-*Cas9-GFP* mice injected with CRISPR AAV targeting the indicated gene or non-targeting control sgRNA (sgCtrl). Two independent sgRNAs were used. **B,** Representative H&E images showing multifocal PanINs in sgCtrl transduced pancreas and PADC tumors in sgUsp15 transduced pancreas. Scale bar 100μm. **C,** Western blot analysis showing loss of protein expression after CRISPR-mediated knockout of Usp15 (two independent sgRNA) in KC cells. **D,** Representative H&E images of mice with the indicated genotype showing multifocal PanINs and PADC tumors. Scale bar 100μm. **E,** Cell proliferation curves of KC cells transduced with the indicated sgRNA obtained using the IncuCyte live-cell imaging and data are expressed as cell confluence percentage (% mean□±□SD, n□=□2). **F,** Cell proliferation curves of KC cells expressing ubiquitin variants inhibiting Usp15 (Ubv15.1a and Ubv15.1/d) or wildtype ubiquitin (Ub^wt^) as control. **G,** Tumor-free survival after orthotopic injection sgCtrl or sgUsp15 KC cells. **H,** Dose-response curves for KPC sgCtrl or sgUsp15 cells treated with the indicated concentration of Olaparib in cell proliferation assay (mean□±□SD). P ≤ 0.05 (*), P ≤ 0.01(**), P ≤ 0.001 (***), P ≤ 0.0001 (****)

Pancreatitis is one of the highest risk factors for the development of PDAC in humans and cooperates with oncogenic *KRas* mutations to induce PDAC formation in *mice^24,28^*. Therefore, we repeated our screen and treated mice with chronic, low doses of cerulein to induce mild pancreatits. As expected, cerulein treatment significantly accelerated PDAC development in KC mice transduced with the PDAC sgRNA library (17 versus 32 weeks median survival, p<0.0001), and a trend towards faster PDAC development in KC mice transduced with the control library (**Supplementary Data Fig. 2e**). In line with the previous screen, *Cdkn2a* was the top-scoring genes followed by *Rnf43* and the newly identified genes, *Usp15* and *Scaf1* (**Supplementary Data Fig. 2f**), further supporting their function as strong suppressors of pancreatic cancer.

### Usp15 is a haploinsufficient PDAC tumor suppressor regulating TGFβ, WNT and NFκb signaling

The multi-domain deubiquitinase USP15 regulates diverse processes such as the p53 tumor suppressor pathway^29^, MAPK signaling^30^, Wnt/beta-catenin signaling^31^, TGF-β signaling^32–34^, NfKb signaling^33,35,36^ and chromosome integrity^37,38^ either through regulated de-ubiquitination of direct substrates such as MDM2, APC, SMADs or TGF-β receptors or de-ubiquitination-independent functions such as through proteinprotein interactions^38^. Interestingly, focal *USP15* copy-number losses have been identified in ~25% of pancreatic cancer cases^39,40^, which was confirmed in an independent large scale genome study^41^ (**Supplementary Data Fig. 3a**).

To validate the tumor suppressive function of Usp15, we first injected KC mice individually with one library or one newly designed sgRNA. All transduced mice developed highly proliferative pancreatic tumors with much shorter latencies compared to mice transduced with the non-targeting control sgRNAs (sgCrtl) (**Fig. 2a**). In fact, age-matched control KC mice only exhibited PanINs at the time when USP15 knockout mice exhibit aggressive PDACs (**Fig. 2b**). All tested tumors harbored bi-allelic frame-shift mutations in the target gene, and western blot analysis confirmed loss of *Usp15* expression (**Fig. 2c** and **Supplementary Data Fig. 3b-d)**.

To further confirm the tumor suppressive role and rule out any confounding effect of the Cas9 endonuclease expression, we generated conditional *Usp15*^fl/fl^; *KRas^G12D^*; Pdx1-Cre. This conventional knockout approach recapitulated our CRISPR/Cas9 findings (**Fig. 2d**), validating our *in vivo* CRISPR approach. Interestingly, *Usp15*^fl/+^ heterozygous mice also presented with significantly shorter disease-free survival (**Fig. 2d**), indicating *Usp15* functions as haploinsufficient tumor suppressor.

Next, we established primary PDAC cell lines from KC mice as well as KC mice with concomitant expression of the hotspot p53^R270H^ mutant (KPC) and used CRISPR/Cas9 to knock-out *Usp15* (**Fig. 2c** and **Supplementary Data Fig. 3d)**. Loss of *Usp15* significantly increased proliferation of these KC cells, while it did not affect KPC cells, presumably, because those cells are at the maximal proliferation rate (**Fig. 2e** and **Supplementary Data Fig. 3e**). Similar results were obtained using ubiquitin variants (UbVs) that bind and block the catalytic domain of Usp15^42^, indicating that this tumor suppressive function is de-ubiquitination dependent (**Fig. 2f**). Upon orthotopic injection, *Usp15* knock-out KC cells also formed allograft tumor faster than non-targeting control cells (**Fig. 2g**). Together, these data show that Usp15 regulates tumor cell proliferation in a cell-autonomous manner and loss of *Usp15* increases a cell’s ability to form allograft tumors.

In line with a previous report^37^, we also found that loss of *Usp15* sensitizes pancreatic cancer cells to Poly-(ADP-ribose) polymerase inhibition (PARPi) by Olaparib. This increased drug sensitivity was stronger in KPC cells than KC cells and was also seen in response to Gemcitabine, one of the most commonly used chemotherapies to treat pancreatic cancer (**Fig. 2h** and **Supplementary Data Fig. 4a-c**). In addition, we found that Olaparib and Gemcitabine treatment significantly increases expression of Usp15 in KC and KPC cells (**Supplementary Data Fig. 4d**). As such, USP15 appears to function as a double-edged sword in pancreatic cancer, where loss of Usp15 enhances tumor progression in the initial stages of tumorigenesis but sensitizes to certain treatment regimens in the later stages.

Given the wide range of USP15 substrates and USP15-regulated pathways with well-known functions in cancer, we set out to elucidate USP15’s exact role in PDAC suppression. First, we transcriptionally profiled primary KC cells transduced with sgRNAs targeting *Usp15* or non-template controls sgRNAs. Inactivation of *Usp15* resulted in dramatic changes in gene expression when compared to scrambled control *Kras^G12D^* tumor cells (794 differentially expressed genes (DEG), false discovery rate (FDR) < 0.05 and absolute log2 fold-change > 1, **Fig. 3a** and **Supplemental Table 3**). Gene set enrichment analyses (GSEA) revealed significantly upregulated gene sets associated with xenobiotic detoxification, glutathione metabolism, anabolic processes, and oxidative phosphorylation (**Fig. 3b** and **Supplemental Table 3**). These findings are in line with USP15’s known role in negatively regulating NRF2 (encoded by the NFE2L2 gene), the master regulator of glutathione metabolism and the redox balance of a cell. In addition, NRF2 expression is induced by oncogenic KRAS and known to stimulate proliferation and suppress senescence of PDAC cells^43^. GSEA also revealed decreased genes sets associated with inflammatory responses, TNFα signaling, TGFβ signalling, and p53 signaling (**Fig. 3b-d and Supplementary Data Fig. 4e**), all pathways with well-known tumor suppressive function in PDAC development^18,44^. Together, these data indicate that Usp15 functions as a strong haploinsufficient PDAC tumor suppressor potentially by regulating the TGFβ signalling pathway.

**Fig. 3.**
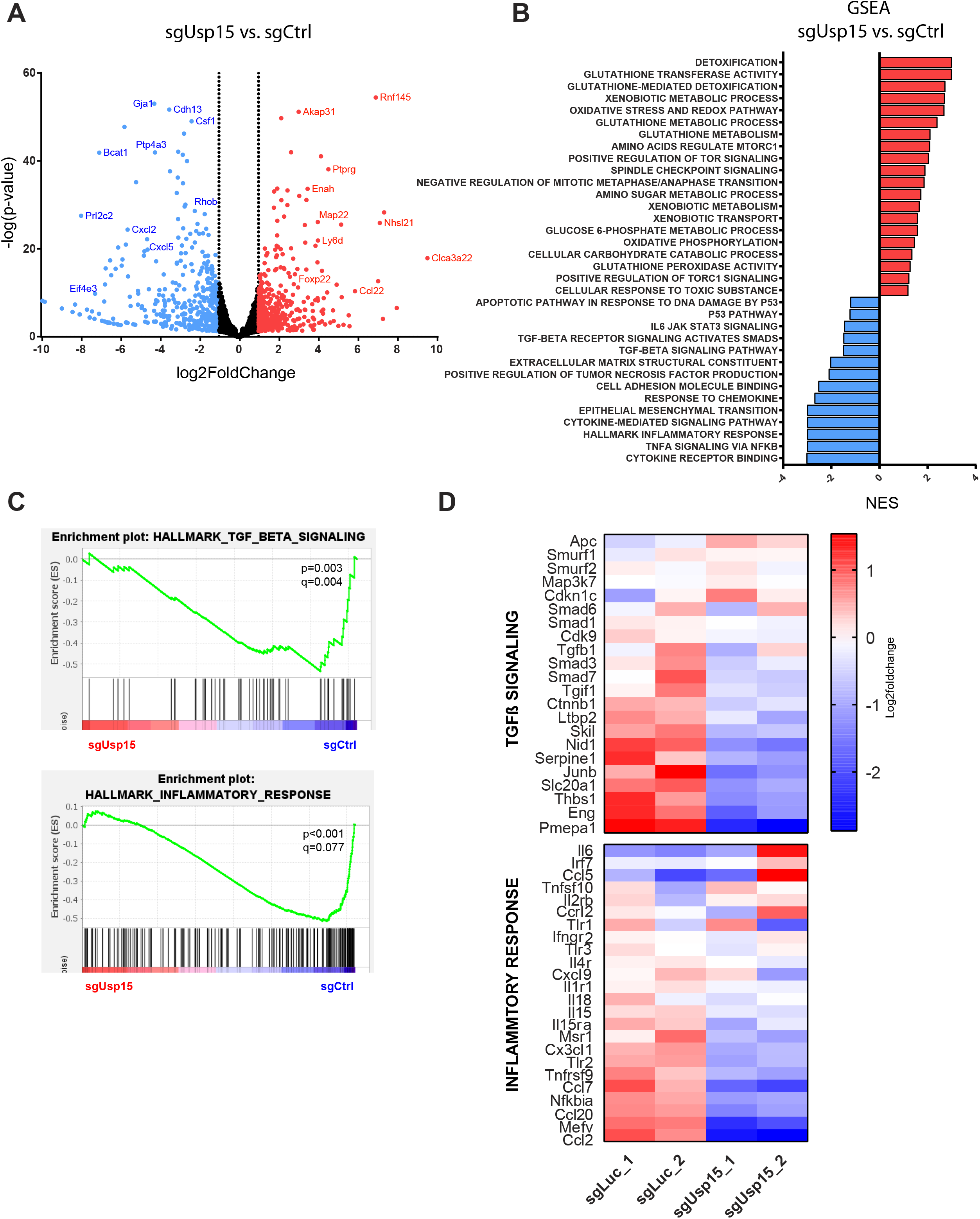
Usp15 regulates several pathways involved in PDAC development. **A,** Volcano Blot showing differential expressed genes between Usp15-knockout compare to control KC cells. **B,** Bar graph showing Gene set enrichment analysis (GSEA) for Usp15-knockout compare to control KC cells, demonstrating strong association with mitotic cell cycle, inflammatory responses, TNFα signaling, TGFβ signaling, and p53 signaling **C,** GSEA plot for Hallmark TGFβ signalling. **D,** Heatmaps of log2 counts per million for selected differentially expressed genes in KC cells.

### SCAF1 is a PDAC tumor suppressor and regulates USP15 levels

Our second new hit, SCAF1 (SR-Related CTD Associated Factor 1), is a member of the human SR (Ser/Arg-rich) superfamily of pre-mRNA splicing factors. It interacts with the CTD domain of the RNA polymerase II (RNAPII) and is thought to be involved in pre-mRNA splicing^45^. Its close homologs SCAF4 and SCAF8 were recently shown to be essential for correct polyA site selection and RNAPII transcriptional termination in human cells^46^. SCAF1 was also one of the top-scoring hits in a screen that looked for genes that can restore homologous recombination in *BRCA1*–deficient cells and thus conferred resistance to PARP inhibition^47^. However, the molecular function of SCAF1 remains completely elusive.

First, we validated the tumor suppressive function of Scaf1 by injecting KC mice individually with one library or one newly designed sgRNA. All transduced mice developed highly proliferative pancreatic cancer with much shorter latencies compared to mice transduced with the non-targeting control sgRNAs (**Fig. 4a, b** and **Supplementary Data Fig. 5a, b**). Similar to *Usp15* knockout cells, we also found that primary *Scaf1* knockout KC cells exhibited increased proliferation in culture and formed tumors faster when injected orthotopically into mice compared to scrambled control KC cells (**Fig. 4c and d**). *Scaf1* knockout cells also exhibited significantly increased sensitivity to Olaparib (**Fig. 4e** and **Supplementary Data Fig. 5c**), again phenocopying *Usp15* knockout cells.

**Fig. 4.**
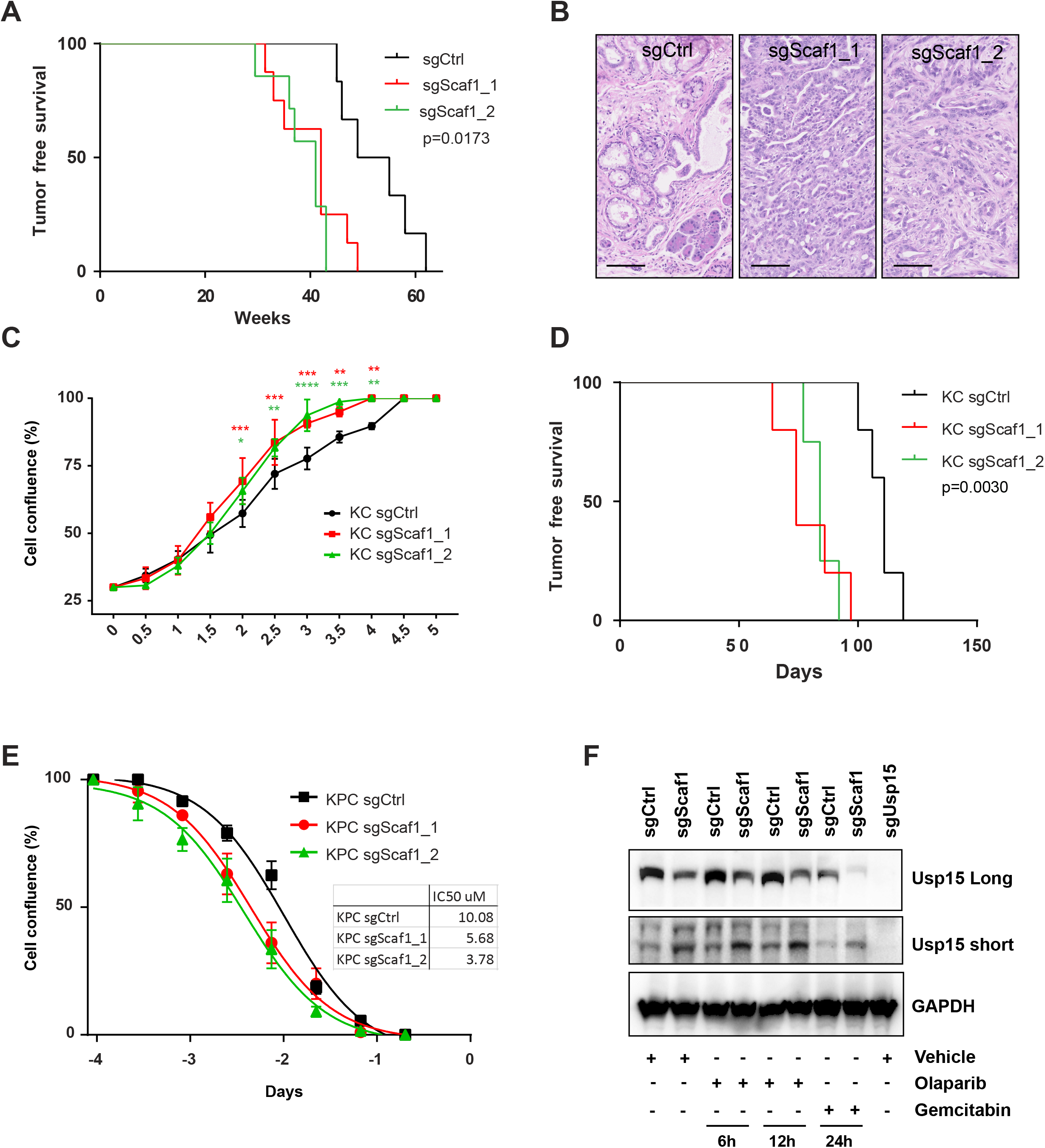
Scaf1 functions as PDAC tumor suppressor. **A,** Tumor-free survival of Pdx1-Cre;LSL-*Kras*^G12D^;LSL-*Cas9-GFP* mice injected with CRISPR AAV targeting the indicated gene or non-targeting control sgRNA (sgCtrl). Two independent sgRNAs were used. **B,** Representative H&E images showing multifocal PanINs in sgCtrl-transduced pancreas and PADC tumors in sgScaf1-transduced pancreas. Scale bar 100μm. **C,** Cell proliferation curves of KC sgCtrl and sgScaf1 cells were obtained using the IncuCyte live-cell imaging and data are expressed as cell confluence percentage (% mean□±□SD). P ≤ 0.05 (*), P ≤ 0.01(**), P ≤ 0.001 (***), P ≤ 0.0001 (****) **D,** Tumor-free survival after orthotopic injection sgCtrl or sgScaf1 KC cells. **E,** Dose-response curves for KPC sgCtrl or sgScaf1 cells treated with the indicated concentration of Olaparib in cell proliferation assay (mean□±□SD). **F,** Western Blot analysis of Usp15 in KC cells transduced with the indicated sgRNAs and incubated with the listed treatment. P ≤ 0.05 (*), P ≤ 0.01(**), P ≤ 0.001 (***), P ≤ 0.0001 (****)

Interestingly, we found a potential connection between Scaf1 and Usp15. *Scaf1* knockout cells exhibited reduced expression of full-length Usp15 (molecular weight of ~125kDa) and showed expression of a 25kDa short Usp15 isoform (**Fig. 4f**). Expression of this short isoform also appears upon treating PDAC cells with Olaparib or Gemcitabine and was also observed in human Panc-1 cells (**Supplementary Data Fig. 4c and 5d**), indicating that this short Usp15 is evolutionary conserved.

To further examine a potential function of this truncated isoform, we cloned and transduced the long and the short isoforms into primary Usp15 knock-out KC cells (**Supplementary Data Fig. 5e**). While fulllength Usp15 was able to supress the hyperproliferative phenotype of Usp15 knock-out cells, the short isoform failed to suppress the cell proliferation (**Supplementary Data Fig. 5f**). Similarly, re-expressing the full-length but not the short Usp15 isoform reversed the sensitivity of Usp15 knock-out KC cells to Olaparib and gemcitabine. In addition, overexpression of the full-length or the short Usp15 isoform did not alter proliferation of wildtype KC cells (**Supplementary Data Fig. 5f**), indicating that the short isoform does not exhibit dominant negative functions. Together this data indicate that the short isoform has no tumor suppressive functions or alters response to PARP inhibition and that the effects of Scaf1 on Usp15 might be limited to the reduced expression of full-length Usp15.

To further elucidate the effects of Scaf1, we transcriptionally profiled Scaf1 knockout KC cells. Inactivation of *Scaf1* resulted in 625 differentially expressed genes (DEG) (false discovery rate (FDR) < 0.05 and absolute log2 fold-change > 1, **Fig. 5a** and **Supplemental Table 3**) when compared to scrambled control *Kras^G12D^* tumor cells. GSEA revealed significantly upregulated gene sets associated with nucleotide metabolism, glutathione metabolism, microtubule polymerization and oxidative phosphorylation as well as downregulates of gene sets associated with TNFα signaling, one-carbon metabolism, xenobiotic catabolic processes, mTorc1/mTOR signaling, hypoxia and p53 signaling. In addition, we found a trend towards downregulated TGFβ signalling (**Fig. 5b** and **Supplemental Table 3**).

**Fig. 5.**
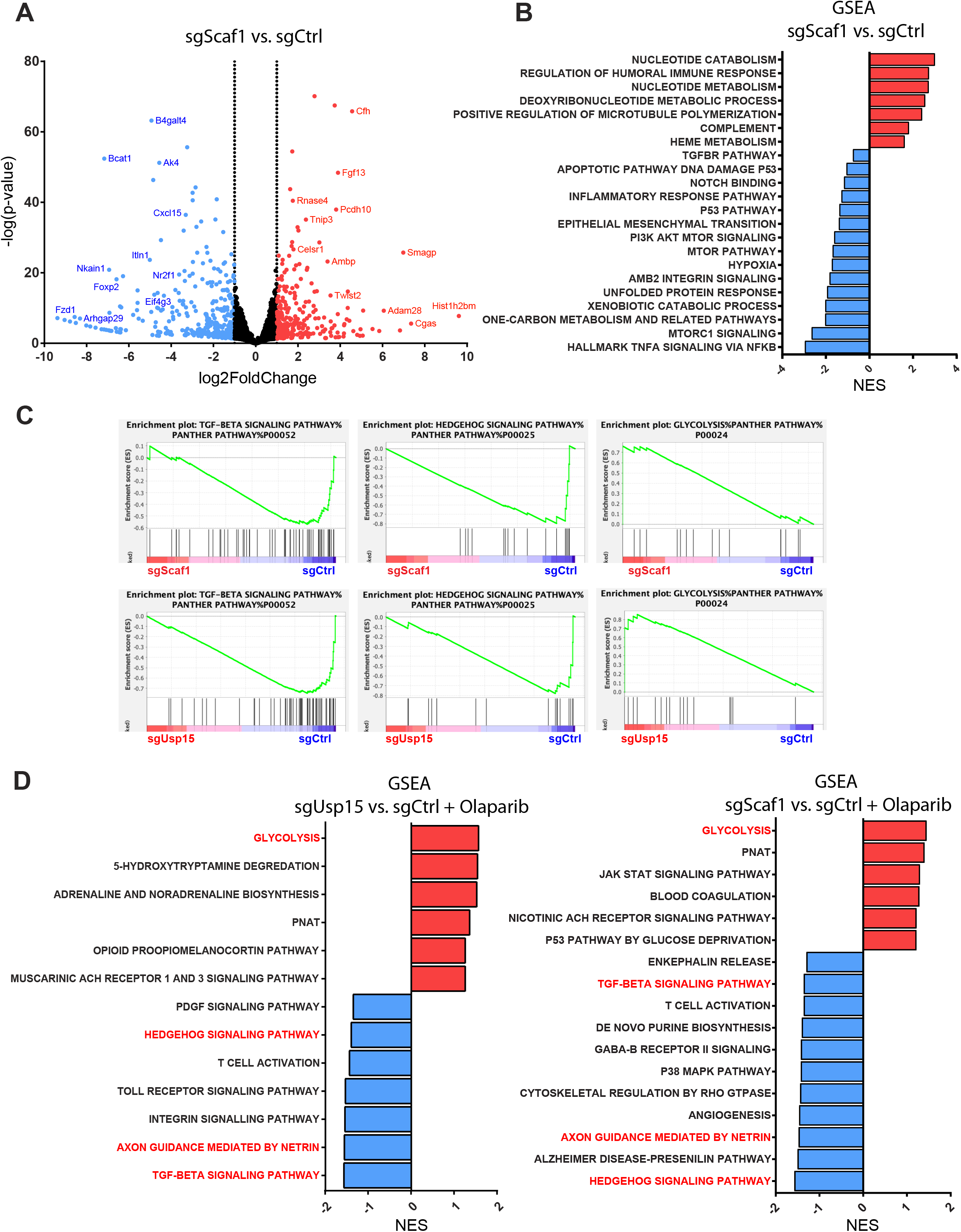
Scaf1 regulates several pathways involved in PDAC development and Olaparib response. **A,** Volcano Blot showing differential expressed genes between Scaf1-knockout compare to control KC cells. **B,** Bar graph showing Gene set enrichment analysis for Scaf1-knockout compare to control KC cells demonstrating strong association with nucleotide metabolism, TNFα signaling, mTorc1/mTOR signaling, hypoxia and p53 signaling. **C,** GSEA plot for Scaf1-knockout and Usp15-knockout compare to control KC cells showing downregulation of hedgehog signalling, TGFβ signalling and upregulation of glycolysis as the top dysregulated pathways upon Olaparib Treatment. **D,** Bar graph showing Gene set enrichment analysis for Scaf1-knockout and Usp15-knockout compare to control KC cells upon Olaparib treatment.

Lastly, we set out to elucidate how Usp15 and Scaf1 regulate the response of pancreatic cancer cells to PARP inhibition. Interestingly, transcriptional profiling and GSEA following Olaparib-treatment revealed that both, Usp15 and Scaf1 knock out cells, exhibited downregulation of hedgehog signalling, TGFβ signalling and ‘axon guidance by netrin’ as well as upregulation of ‘glycolysis’ as the top dysregulated pathways compared to Olaparib-treated control KC cells (**Fig. 5c** and **Supplemental Table 4**). Together, this indicates a common mechanism leading to increased sensitivity to PARP inhibition shared between Usp15 and Scaf1 knock out cells. Thus, Scaf1 and Usp15 knockout cells share several alterations such as upregulated TNFα signaling and downregulated TGFβ, hedgehog and p53signalling but also several distinct pathways.

## Discussion

One key bottleneck on the path towards ‘Precision Medicine’ is our fragmentary understanding of the functional consequence of most genetic alterations associated with specific malignancies. Cancer develops due to the acquisition of cooperating alterations in tumor suppressor and oncogenes (=driver mutations), which are thought to either occur gradually or simultaneously in a single catastrophic event (e.g. chromothripsis) as recently shown by Notta *et al*.^48^. Through these mutational processes tumors also accumulate hundreds of random bystander mutations, which make it exceedingly hard to interpret genomic data and identify the few real driver mutations that trigger tumor initiation, progression, metastasis and therapy resistance. Even within known cancer genes, many variants are of uncertain significance (VUS), where the effect of the genetic alteration on gene function cannot be predicted using current bioinformatics tools. Genetic-based treatment design is thus reliant on weeding out bystanders and identifying bona fide driver mutations, as only the latter have diagnostic and therapeutic value. Secondly, we have to identify the actionable nodes within a given cancer gene network that can be exploited to selectively kill or disable cancer cells. Thirdly, we have to identify cancer genotypes that are sensitive to a given treatment and those genotypes that confer resistance to be able to stratify patients into the best treatment arm. Lastly, we have to establish efficient animal models to test the efficacy of novel therapeutic strategies and anticipate and overcome resistance mechanisms.

Our *in vivo* PDAC CRISPR/Cas9-screen identified several novel *bona fide* PDAC tumor suppressor genes such as *USP15* and *SCAF1*. *USP15* is a broadly expressed deubiquitinase and was implicated in several cancer associated pathways. For example, USP15 can act as a tumor promoter in estrogen receptor positive breast cancer by deubiquitinating and thereby stabilizing the estrogen receptor^49^, by stabilizing TGF-β receptor 1 (TGFβR1) in glioblastoma^32^, or by deubiquitinating and stabilizing MDM2, leading to p53 inactivation^29^. USP15 was also shown to play important roles in inflammation in response to an infectious and autoimmune insults and following tissue damage^50^. In line with these reports, we found that loss of USP15 in pancreatic epithelium leads to reduced TGF-β signaling as well as downregulation of inflammatory responses to cytokine and chemokines such as TNFa and IL6 signaling.

In pancreas cancer cell lines, Peng et al. showed that USP15 regulates homologous recombination and DNA double strand break (DSB) repair by deubiquitinating BARD1, thereby promoting BARD1-HP1γ interaction and increased BARD1-BRCA1 retention at DSB. Mutation or loss of USP15 impairs DSB repair and thus leads to increased sensitivity to PARP inhibition^37^. We recapitulated this data and also showed increased sensitivity to PARP inhibition but also increased sensitivity to Gemcitabine, the most common PDAC chemotherapy. The increased sensitivity to Gemcitabine was surprising at first, as Gemcitabine does not induce DNA DSB. Transcriptional profiling of *USP15* knock-out cells showed that glutathione metabolism and oxidative stress and redox pathways are significantly upregulated, indicating that *USP15* knock-out cells are experiencing increased cellular stress. This could conceivably further explain the increased sensitivity to Olaparib but also to Gemcitabine. In addition, we found that In addition to USP15’s role in regulating sensitivity to PARP inhibition, we now found that USP15 itself functions as a strong tumor suppressor in pancreatic cancer. Importantly, our data indicates that USP15 functions as a haploinsufficient tumor suppressor in pancreas cancer. The growing list of haploinsufficient cancer driver genes identified in genetic screens^11,51–56^ raises the provocative question whether the lack of comprehensive screening within innate tumor microenvironment obstructed our capabilities of identifying many of these haploinsufficient cancer driver genes. This is in line with recent findings from Martin et al., showing that the adaptive immune system is a major driver of selection for tumor suppressor gene inactivation^57^. Historically, most attention has focused on frequently mutated dominant oncogenes and recessive tumor suppressor genes, but recent large scale genomic efforts revealed recurrent copy number alterations (CNA) mainly involving shallow losses or gains of large regions^58,59^. The remarkably recurrent, specific pattern of these CNAs certainly indicates that one or several genes in these regions are being selected for, presumably by loss of haploinsufficient tumor suppressor genes^60^. However, with a few exceptions ^61–64^, cancer driver genes conferring selective advantage of certain CNA are virtually unknown. Bioinformatic approaches to delineate cancer driver from passenger mutations are usually based on statistical enrichment of specific patterns of somatic point mutations and/or amino acid conservation signifying functional importance. However, CNA are simply too large sometimes spanning hundreds-to-thousands of genes, too numerous and too noisy and most studies are underpowered to call driver genes by bioinformatic means. Given that some CNAs are linked to worse outcome and might have therapeutic implications, functional annotating recurrent haploinsufficient cancer driver genes in recurrent CNAs is of high clinical relevance. Importantly, it would be of great significance to test whether PDAC patients with *USP15* or *SCAF1* losses indeed show increased sensitivity to Gemcitabine or Olaparib and exhibit a better therapeutic outcome, as observed with our findings. Together, this study highlights the utility of *in vivo* CRISPR screening to integrate cancer genomics and mouse modeling for rapid discovery, validation and characterization of novel PDAC genes.

## Data Availability

All RNA-seq are will be made available at NCBI Gene Expression Omnibus.

## Acknowledgements

We thank all members of our laboratories for helpful comments, with additional thanks to Y.Q. Lu, and G. Mbamalu for their insight and assistance. We also thank The Centre for Phenogenomics, Network Biology Collaborative Centre and Flow Cytometry facility at LTRI as well as the Flow Cytometry Facility at the University of Toronto. Funding: This work was supported by an OICR Translational Research Grant to D.S., S.G. and F.N. and a Canadian Institute of Health project grant to K.C. and D.S (PJT175270). The development of *Usp15^flox/flox^* mice was supported in part by a consortium grant from the Healthy Brains for Healthy Lives McGill program, the Consortium Quebecois de la Recherche sur le Médicament, Brain Canada, and Corbin Therapeutics.

## Author Contributions

S.M. performed all experiments. A.M. performed bioinformatics analysis. T.W and F.N performed the PDO experiments, M.G. and K.C. analysed the RNAseq experiments. D.D., K.N.A. and R.T. helped with mouse and RT-PCR experiments. P.G. and S.S.S provided the Usp15^fl/fl^ and the UbV variants, respectively and helped with experimental design. R.W. and A.S performed and analysed the ribosome profiling. D.S. coordinated the project and the experiments and together with S.M. wrote the manuscript.

## Competing interests

All authors declare no competing interests.

## Supplementary Materials

Materials and Methods

Figures S1-S5

Tables S1-S4

References 1 – 68

## Supplementary Data Figure legends

**Supplementary Fig. 1. *In vivo* CIRPSR knock out efficiency in murine pancreas. A**, Image of AAV injection into the pancreas. Representative immunohistochemistry of pancreas injected with control or AAV H2B-RFP. **B,** Representative immunofluorescence of Pdx1-Cre; LSL-KRasG12D epithelial cells transduced with AAV H2B-RFP. Scale bar 50 μm. **C,** Representative images showing GFP and H2B-RFP expression in Pdx1-Cre R26-LSL-Cas9-GFP; transduced with sgGFP-H2B-RFP AAV or control non-targeting sgCTRL-H2B-RFP AAV. Flow cytometry analysis shows the percent of GFP+/H2B-RFP+ double cells. **D** Representative image of Pdx1-Cre; LSL-KRasG12D; R26-LSL-Cas9-GFP pancreas injected with sgTp53. Tumor-free survival of PDX1-Cre LSL-KRasG12D LSL-Cas9-GFP mice transduced with a sgTp53 or sgCtrl. **E,** Representative images of pancreas transduced with an AAV-GFP/AAV-RFP mixture showing cells transduced with GFP or RFP or double-infected expressing GFP+/RFP+ cells. More double positive cells are observed at higher viral titre. **F,** Representative, image of reporter LSL-KRasG12D R26-LSL-Confetti pancreas infected with sgRNA-Cre AAV.

**Supplementary Fig. 2. *In vivo* pancreatic cancer CRISPR screen A.** Graph showing sgRNA correlation and representation for PDAC and CTRL libraries in plasmid DNA versus infected MEFs DNA. Each dot represents a guide. **B.** Representative image of H2B-RFP liver and lung metastasis. Representative H&E showing a PDAC-Library liver and lung metastasis Scale bar 100 μm. Representative immunofluorescence of PDAC-Library liver metastasis showing H2B-RFP and CK19 expression. Scale bar 50 μm. **C.** Percentage of Pdx1-Cre LSL-*KRas*^G12D^ LSL-*Cas9-GFP* mice transduced with corresponding sgRNA libraries displaying metastasis. **D.** Representative pie charts showing tumor suppressor genes with enriched sgRNAs in tumor DNA obtained from matching pancreatic tumor, liver and lung metastasis. **E.** Tumor-free survival of Pdx1-Cre LSL-*KRas*^G12D^ LSL-*Cas9-GFP* mice transduced with a PDAC or CTRL library and treated with cerulein. **F.** Column bar graph showing putative tumor suppressor genes with enriched sgRNAs in tumor DNA obtained from PDAC mouse model (sgRNA enriched per tumors are denoted in color).

**Supplementary Fig. 3.**

**A.** Oncoprint of the indicated genes identified as tumor suppressors in our screen as well as alterations in KRas in PDAC Samples (n= 293) **B.** Representative sanger sequencing plots showing discordance of DNA from a sgUsp15-targeted sample compared to a control sample. **C.** Gene editing efficiency os sgUsp15. Efficiency was determined using sanger-sequencing data of PCR-amplified sgRNA target sites followed by Tracking of Indels by Decomposition (TIDE https://tide.nki.nl) algorithm on PDAC cells. **D.** Western blot analysis showing loss of protein expression after CRISPR-mediated knockout of Usp15 (two independent sgRNA in KPC cells. **E.** Cell growth curves of KPC cells sgCtrl and sgUsp15. Data are expressed as cell confluence percentage (%; mean□±□SD, n□=□2).

**Supplementary Fig. 4.**

**A.** Dose-response curves for KC sgCtrl or sgUsp15 cells treated with the indicated concentration of Olaparib in cell proliferation assay (%; mean□±□SD, n□=□2). **B.** Dose-response curves for KPC and KC sgCtrl or sgUsp15 cells treated with the indicated concentration of Gemcitabine in cell proliferation assay (%; mean□±□SD, n□=□2). **C.** Cell response for KC expressing different ubiquitin variants treated with 4μM of Olaparib in cell survival assay (%; mean□±□SD, n□=□2). Cell percentage normalized to control. **D.** Western blot analysis showing USP15 isoforms expression in KC cells transduced with AAV-sgCtrl. Cell were treated with the specified drug for the listed duration. **E.** GSEA enrichment plots of differentially expressed pathways associated with loss of Usp15.

**Supplementary Fig. 5.**

**A.** Gene editing efficiency of sgScaf1. **B.** RT-PCR analysis of Scaf1 expression in KPC and KC cells transduced with two independent sgRNA. **C.** Dose-response curves for KC sgCtrl or sgScaf1 cells treated with the indicated concentration of Olaparib in cell proliferation assay (%; mean□±□SD, n□=□2). **D.** Western blot analysis showing USP15 isoforms expression after CRISPR-mediated knockout of Scaf1 in KPC cells transduced with AAV-sgCtrl, AAV-sgUsp15 or AAV-sgScaf1. E. Western blot analysis showing USP15 isoforms construct expression. Schematic illustration of the domain organization of USP15 isoforms. USP15 catalytic domain is shown in red. USP15 active triad C269, H862, and D879 are denoted by yellow lines **F.** Cell growth curves of KC cells sgCtrl and sgUsp15 expressing listed isoforms of USP15. Data are expressed as cell confluence percentage (%; mean□±□SD, n□=□2). G. Dose-response curves for KPC and KC sgCtrl or sgUsp15 cells expressing the expressing listed isoforms of USP15 and treated with the indicated concentration of Gemcitabine and Olaparib in cell proliferation assay (%; mean□±□SD, n□=□2). **H.** Cell growth curves of KC cells expressing listed isoforms of USP15. Data are expressed as cell confluence percentage (%; mean□±□SD, n□=□2)

## Methods

### Animals

Animal husbandry, ethical handling of mice and all animal work were carried out according to guidelines approved by Canadian Council on Animal Care and under protocols approved by the Centre for Phenogenomics Animal Care Committee (18-0272H). The animals used in this study were Pdx1-Cre;KrasG12D/+ mice [B6.FVB-Tg(Pdx1-cre)6Tuv/J;LSL-Kras-G12D] in a mixed C57/Bl6-FVBN background. R26-LSL-Cas9-GFP [#026175 in C57/Bl6 background from Jackson laboratories]. LSL-Kras-G12D; p53-LSL-R270H [B6.129-Krastm4Tyj; 129S4-Trp53tm3Tyj]. LSL-USP15-tm1c [C57BL / 6N-Usp15tm1c(EUCOMM)Wtsi / Tcp] kindly provided by Philippe Gros. CRISPR screens in the Pdx1-Cre;KrasG12D/+; Cas9 cohort were performed in a F1 FVBN/C57Bl6 background. Genotyping was performed by PCR using genomic DNA prepared from mouse ear punches. When total tumor mass per animal exceeded 1000mm3, mice were monitored bi-weekly and scored in accordance to SOP “#AH009 Cancer Endpoints and Tumour Burden Scoring Guidelines”.

### Adeno-associated virus constructs and library construction

sgRNAs targeting pancreatic cancer long tail genes were obtained from Hart *et al.,^65^* (4 sgRNAs/gene) and non-targeting sgRNAs were obtained from Sanjana *et al.,^66^* ordered as a pooled oligo chip (CustomArray Inc., USA) and cloned into AAV sgRNA-H2B-RFP engineered from AAV:ITR-U6-sgRNA(backbone)-pCBh-Cre-WPRE-hGHpA-ITR kindly provided by Feng Zhang (Addgene plasmid #60229) We excluded frequent and known pancreatic cancer tumor suppressor genes such as *TP53* or *Smad4* from the Cancer long tail genes library. The non-targeting sgRNAs are those designed not to target in the mouse genome as negative control. Ad-Cre and Ad-GFP was purchased from the Vector Core at the University of Iowa.

### AAV production and transduction

293AAV cells (AAV-100, Cell biolabs inc) were seeded on a poly-L-lysine coated 15 cm plates and transfected using PEI (polyethyleneimine) method in a non-serum media with AAV construct of interest along with AAV packaging plasmids pAAV-DJ Vector and pHelper Vector. 8 hours post-transfection media was added to the plates supplemented with 10% Fetal bovine serum and 1% Pencillin-Streptomycin antibiotic solution (w/v). 48 hours later, the viral supernatant and cell pellet were collected. Cell lysis was performed by four rounds of freeze/thaw cycles using a dry ice/ethanol bath and filtered through a Stericup-HV PVDF 0.45-μm filter, and then concentrated ~2,000-fold by ultracentrifugation in a MLS-50 rotor (Beckman Coulter). Viral titers were determined by infecting the R26-LSL-tdTomato MEFs and FACS based quantification. *In vivo* viral transduction efficiency was determined by injecting decreasing amounts of a single viral aliquot of known titer, diluted to a constant volume of 10 μl per pancreas. We collected pancreas at 7 days post-infection and determined percent infection using FACS.

### Pancreas viral transduction

Four to six weeks-old mice were anesthetized using 3% isofluorane. Mice were subjected to laparatomy, and injected with 10 uL of purified AAV solution resuspended in PBS using a 28-gauge needle throught the head and tail of the pancreas by slowly retracting the needle. Successful administration was confirmed by a uniform swelling of the injected area. Laparotomies were subsequently closed with a two-layer suture. The majority of the mice survived this operation with no observed complications. Mice were then sacrificed at various days post injection, and pancreatic tissue was harvested for DNA extraction and immunohistochemistry. To verify the sgRNA abundance and representation in the control and pancreas long-tail genes libraries, MEFs were transduced with library virus and collected 48h post transfection. Genomic DNA from all samples was extracted using a QIAamp DNA tissue mini kit (Qiagen). Barcode preamplification, sequencing and data processing were performed as described below. Ad-Cre and Ad-GFP were injected at a final pfu of 1.10^7 pfu/mL.

### Deep Sequencing: sample preparation, pre-amplification and sequence processing

Genomic DNA from epithelial and tumor cells were isolated with the DNeasy Blood & Tissue Kit (Qiagen). 5μg genomic DNA of each tumor was used as template in a pre-amplification reaction using unique barcoded primer combination for each tumor with 20 cycles and Q5 High-Fidelity DNA Polymerase (NEB). The following primers were used:

FW:5’AATGATACGGCGACCACCGAGATCTACAC**<u>TATAGCCT</u>**ACACTCTTTCCCTACACGACGCT CTTCCGATCTtgtggaaaggacgaaaCACCG-3’
RV:5’CAAGCAGAAGACGGCATACGAGAT**<u>CGAGTAAT</u>**GTGACTGGAGTTCAGACGTGTGCTCTT CCGATCTATTTTAACTTGCTATTTCTAGCTCTAAAAC-3’

The underlined bases indicate the Illumina (D501-510 and D701-712) barcode location that were used for multiplexing. PCR products were run on a 2% agarose gel, and a clean ~200bp band was isolated using Zymo Gel DNA Recovery Kit as per manufacturer instructions (Zymoresearch Inc.). Final samples were quantitated then sent for Illumina Next-seq sequencing (1 million reads per tumor) to the sequencing facility at Lunenfeld-Tanenbaum Research Institute (LTRI). Sequenced reads were aligned to sgRNA library using Bowtie version 1.2.2 with options -v 2 and -m 1. sgRNA counts were obtained using MAGeCK count command. A detailed cloning protocol can be found at Loganathan *et al*.,^67^.

### Analysis of genome editing efficiency

LSL-Cas9-GFP MEFs, KPC-LSL-Cas9-GFP and KC-LSL-Cas9-GFP were cultured and infected with AAV carrying corresponding sgRNAs. Cells were live sorted for GFP+/RFP+ expression and expanded further to extract genomic DNA using DNeasy Blood & Tissue Kit (Qiagen). Genomic DNA from tumors from the mice injected with single sgRNAs were also isolated using the same kit. PCR was performed flanking the regions of sgRNA on genomic DNA from both WT cells and cells infected with respective virus or tumors and sent for Sanger sequencing. Gene editing efficiency was determined by Tracking of Indels by Decomposition (TIDE https://tide.nki.nl) algorithm.

### Antibodies

The following primary antibodies were used in this study: Anti-Cytokeratin 19 antibody [RCK108] (1:200, Abcam ab9221), Anti-USP15 monoclonal antibody (M01), clone 1C10 (1:500, Novus Biological H00009958-M01), Anti-GAPDH (6C5) (1:1000, Santa Cruz sc-32233)

### Cell culture

Primary mouse tumor cells KPC and KC were cultured in DMEM supplemented, 10% FBS and Pen Strep. Panc1 cells were cultured in DMEM supplemented, 10% FBS and Pen Strep. Cells were cultured in monolayer for growth and transfection with AAV CRISPR construct containing Cre or H2B-RFP resistance and sgRNA targeting genes of interest. Cells were tested for cutting efficiency post selection with TIDE described earlier and by western blot.

### Immunofluorescence

Tissue sections were fixed with 4% paraformaldehyde for 10 minutes. Following fixation, slides were rinsed 3 times with PBS for 5 minutes. For cells, permeabilization was carried out using 0.5% Tween-20 in PBS at 4°C for 20-minutes and rinsed with 0.05% Tween-20 in PBS for 5 minutes, 3 times each at room temperature. Samples were blocked at room temperature with blocking serum (recipe: 1% BSA, 1% gelatin, 0.25% goat serum 0.25% donkey serum, 0.3% Triton-X 100 in PBS) for 1 hour. Samples were incubated with primary antibody diluted in blocking serum overnight at 4°C followed by 3 washes for 5 minutes in PBS. Secondary antibody was diluted in blocking serum with DAPI and incubated for 1 hour at room temperature in the dark. Following incubation, samples were washed 3 times for 5 minutes in PBS. Coverslips were added on slides using MOWIOL/DABCO based mounting medium and imaged under microscope next day. For quantification, laser power and gain for each channel and antibody combination were set using secondary only control and confirmation with primary positive control and applied to all images.

### RNA-seq and GSEA analyses

RNA was extracted from cells using Quick-RNA plus mini Kit (Zymoresearch Inc., #R1057) as per the manufacturer’s instructions. RNA quality was assessed using an Agilent 2100 Bioanalyzer, with all samples passing the quality threshold of RNA integrity number (RIN) score of >7.5. The library was prepared using an Illumina TrueSeq mRNA sample preparation kit at the LTRI sequencing Facility, and complementary DNA was sequenced on an Illumina Nextseq platform. Sequencing reads were aligned to mouse genome (mm10) using Hisat2 version 2.1.0 and counts were obtained using featureCounts (Subread package version 1.6.3) ^68^. Differential expression was performed using DESeq2 ^69^ release 3.8. Gene set enrichment analysis was performed using GSEA version 4.0; utilizing genesets obtained from MSigDB (https://www.gsea-msigdb.org/gsea/msigdb). For integration with human and existing mouse tumor models, clustering was conducted after normalization and filtering for only intrinsic genes as described previously^77^.

### Statistics and reproducibility

All quantitative data are expressed as the mean□±□SD. Differences between groups were calculated by two-tailed Student’s t-test, Wilcoxon Rank-Sum test (when data was not normally distributed) or Log-rank test for survival data using Prism 7 (GraphPad software).

## Notes

### Competing Interest Statement

The authors have declared no competing interest.

