## Supplementary figures for "*In vivo* CRISPR screens reveal SCAF1 and USP15 as novel drivers of pancreatic cancer"

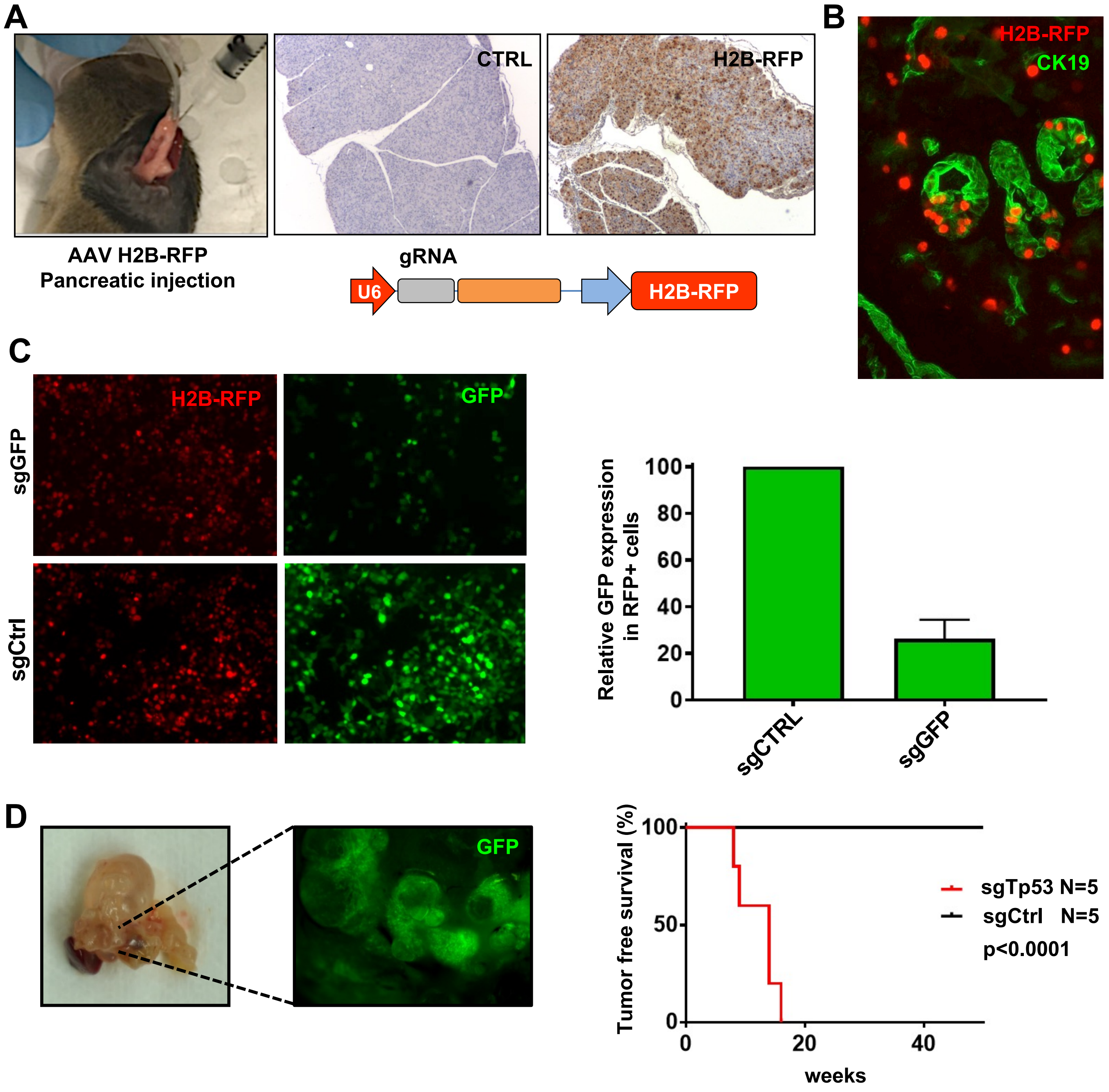


**Supplementary Fig. 1.** ***In vivo*** **CIRPSR knock out efficiency in murine pancreas.**

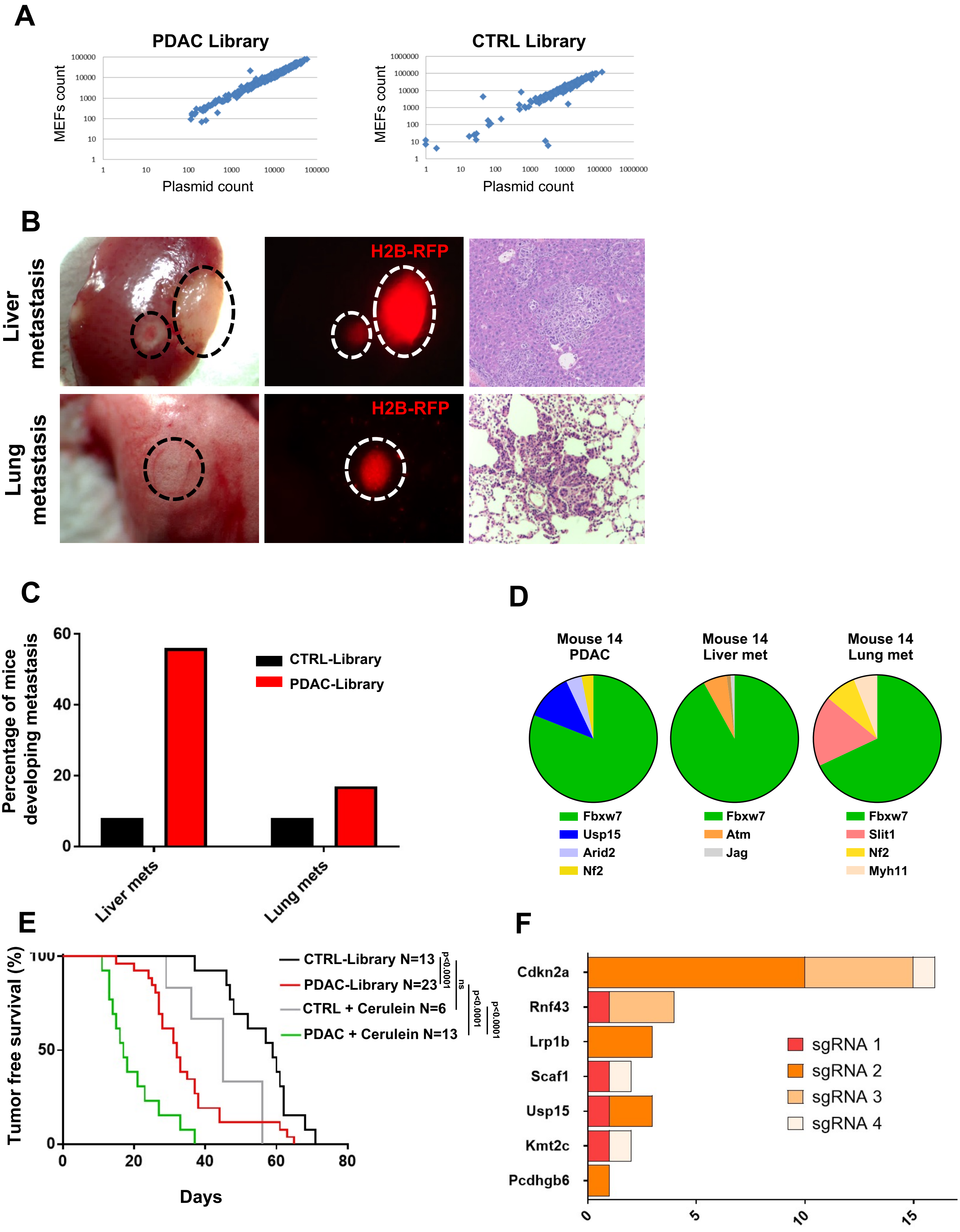


### Extended Figure 2. *In vivo* CRISPR screen

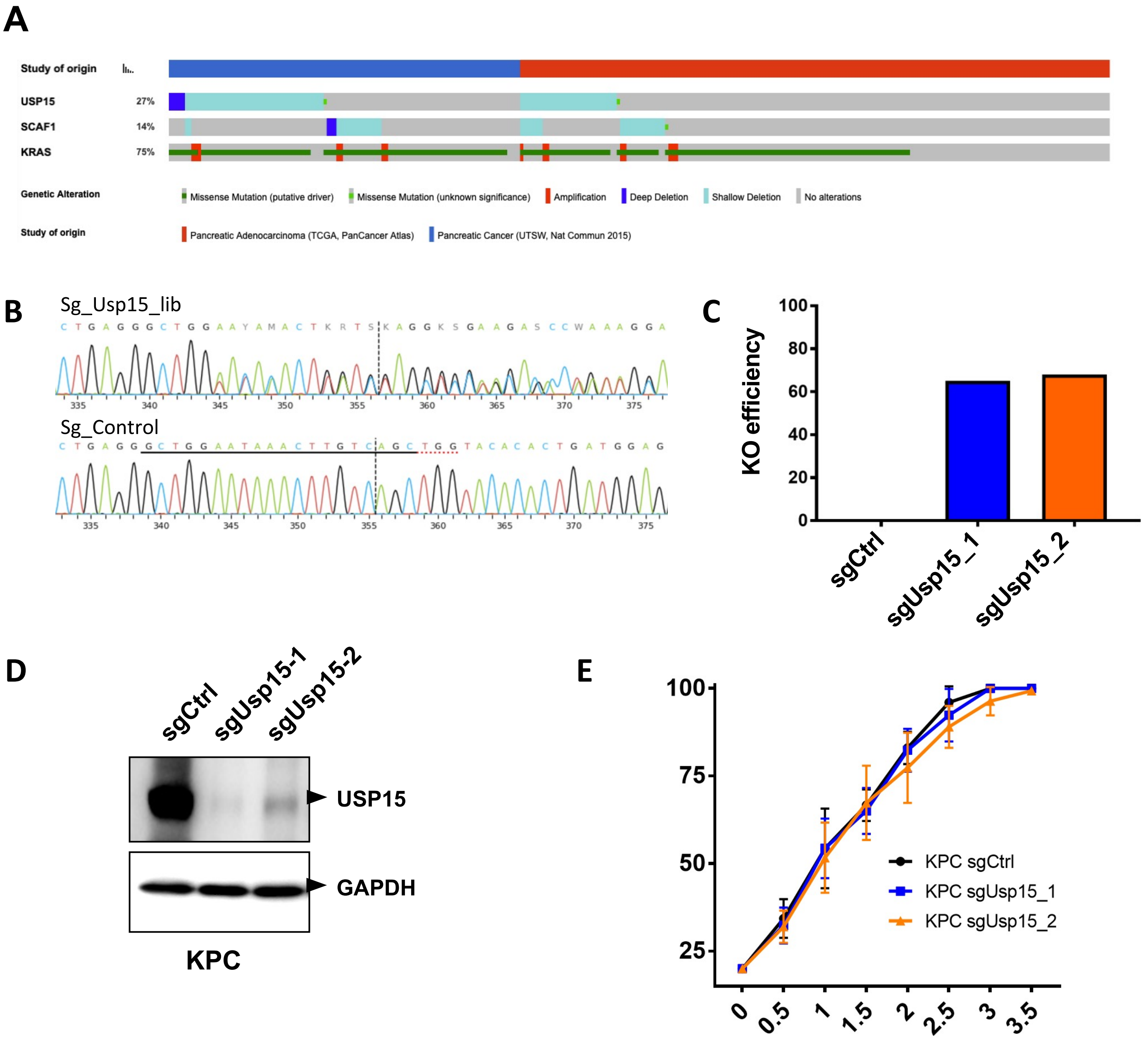


**Extended Figure 3.** ***USP15*** **is a** ***bona-fide*** **PDAC suppressor**

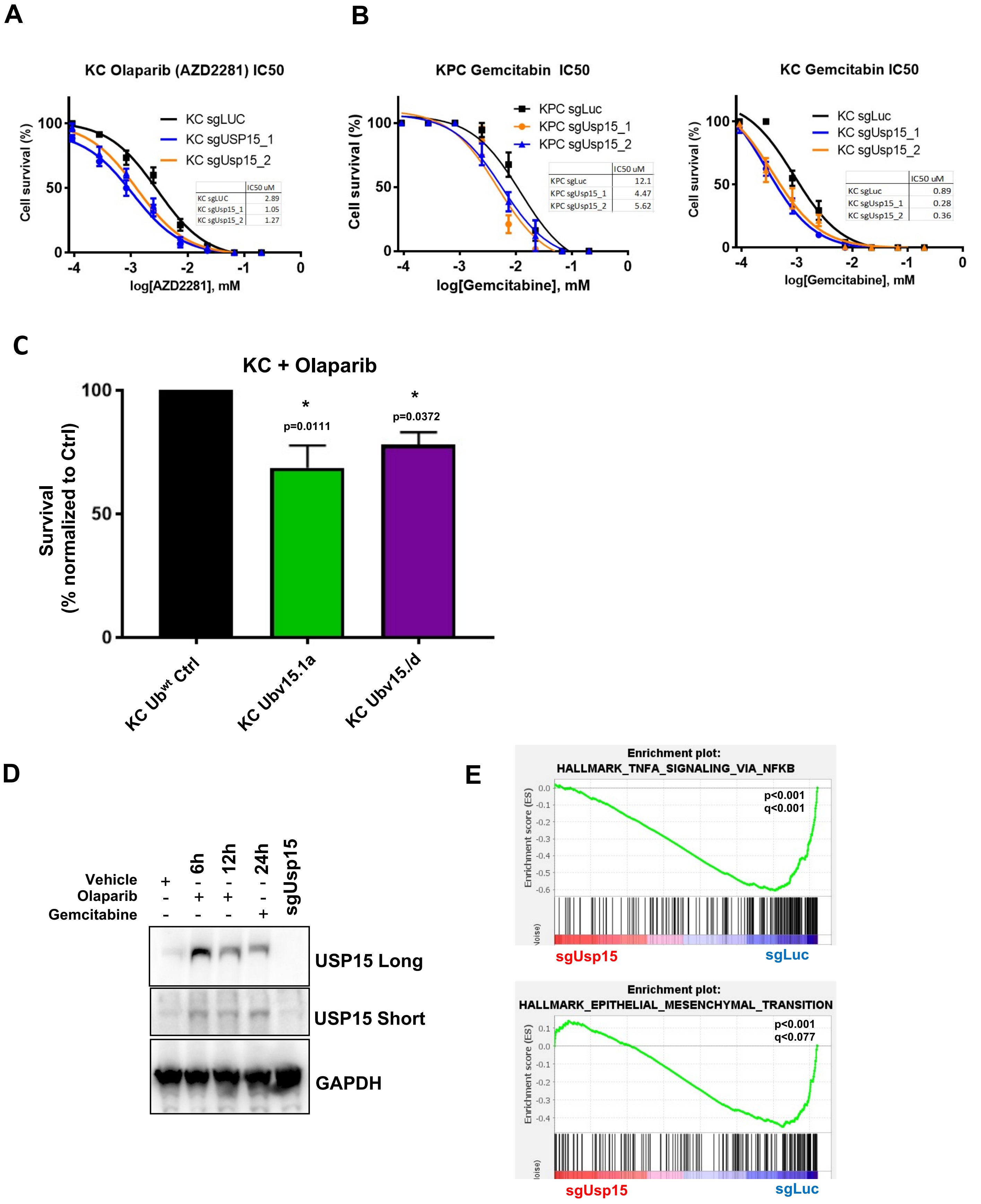


### Extended Figure 4. *USP15* regulates response to PARPi and Gemcitabine

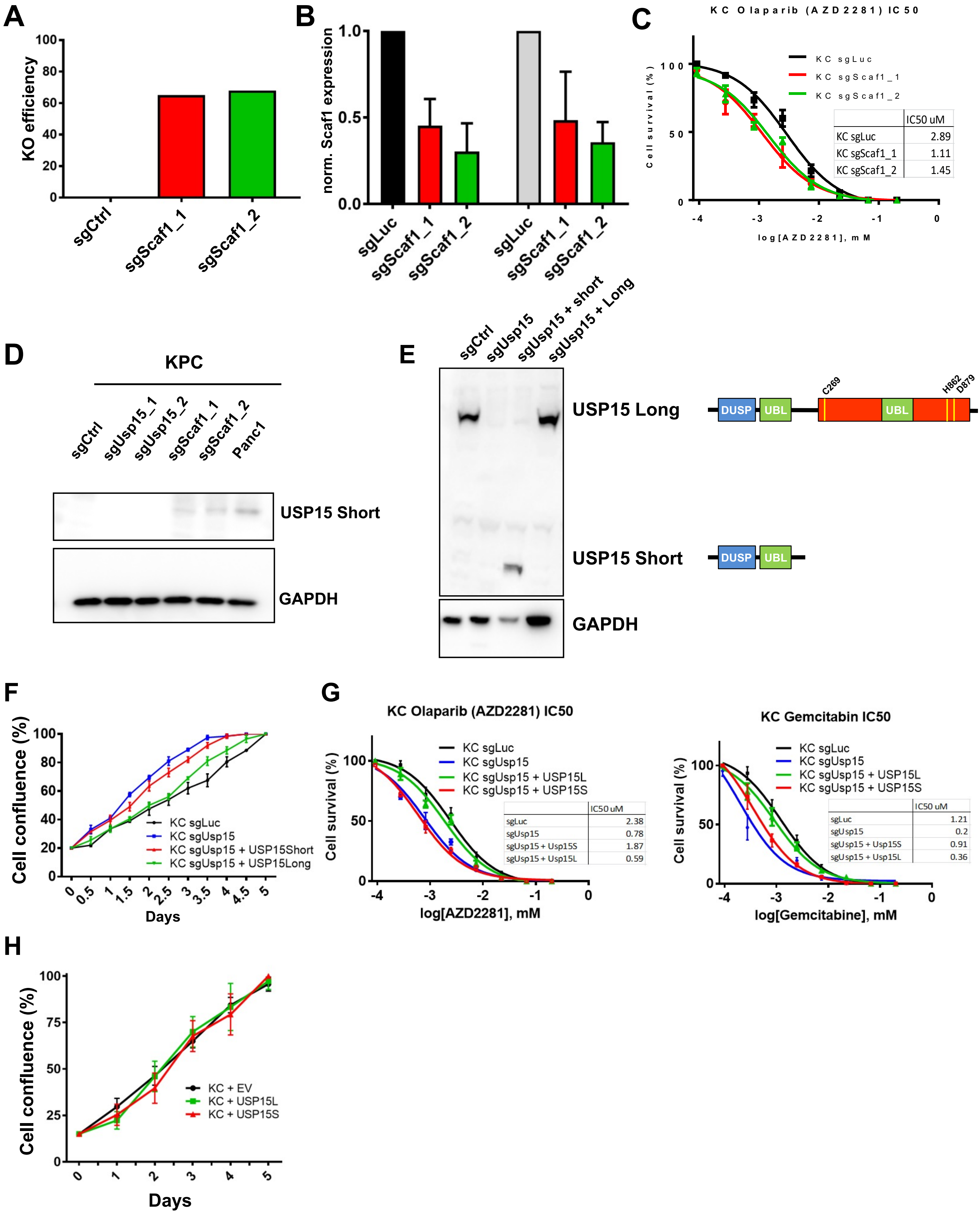


**Extended Figure 5.** ***SCAF1*** **is a** ***bona-fide*** **PDAC suppressor**
